# Evolutionary trajectories of three independent neo-sex chromosomes in *Drosophila*

**DOI:** 10.1101/2021.03.11.435033

**Authors:** Masafumi Nozawa, Yohei Minakuchi, Kazuhiro Satomura, Shu Kondo, Atsushi Toyoda, Koichiro Tamura

## Abstract

Dosage compensation (DC) on the X chromosome is a mechanism to counteract the deleterious effects by gene loss from the Y chromosome. However, DC cannot work efficiently if the X chromosome also degenerates. This indeed occurs in the neo-sex chromosomes in *Drosophila miranda*, where neo-X as well as neo-Y chromosomes are under accelerated pseudogenization. To examine the generality of this pattern, we investigated the evolution of two additional neo-sex chromosomes that independently emerged in *D. albomicans* and *D. americana* and compared their evolutionary processes with that in *D. miranda*. Comparative genomic and transcriptomic analyses revealed that the pseudogenization rate on neo-X is also accelerated in the two species (though lesser extent in *D. americana*). We also found that neo-X-linked genes whose neo-Y homologs are pseudogenized tend to be upregulated more stringently than those whose neo-Y homologs remain functional. Moreover, the genes under strong functional constraints and highly expressed in the testis tended to remain functional on neo-X and neo-Y, respectively. Focusing on the *D. miranda* and *D. albomicans* neo-sex chromosomes that independently emerged from the same autosome, we further found that the same genes tend to have been pseudogenized in parallel on neo-Y. Those genes include *Idgf6* and *JhI-26* whose functions seem to be unnecessary or could be even harmful for males. These results indicate that neo-sex chromosomes in *Drosophila* share a common evolutionary trajectory after their emergence, which may be applicable to other sex chromosomes in a variety of organisms to avoid being an evolutionary dead-end.

## INTRODUCTION

Sex chromosomes are widely present in divergent groups of organisms (Bachtrog et al. 2014). Species with sex chromosomes can maintain a stable sex ratio (i.e., ∼1), irrespective of their surrounding environment, which may be advantageous under certain conditions. However, once sex chromosomes emerge from a pair of autosomes by acquiring sex-determining and sexually antagonistic genes, the proto-X and proto-Y chromosomes mostly stop meiotic recombination to retain stable sex determination, resulting in Y chromosome (hereafter, Y) degeneration (Charlesworth et al. 2005). Consequently, the number of genes on Y is much smaller than that on the X chromosome (hereafter, X) in many species (Koerich et al. 2008; Cortez et al. 2014; Zhou et al. 2014; Dupim et al. 2018; Bracewell and Bachtrog 2020). Therefore, the number of many X-linked genes become imbalanced, i.e., two in females and one in males. Thus, Y degeneration could potentially be deleterious for males and eventually the entire species with sex chromosomes.

Dosage compensation (DC) as a mechanism to counteract this dosage imbalance of X-linked genes between sexes was originally proposed by Muller (1932) and later more clearly by Ohno (1967) (see Gartler 2014). Many researchers have examined DC since then and reported the presence of global or chromosome-wide DC in many organisms (Disteche 2012; Graves 2016) (see also Ellegren et al. 2007; Zha et al. 2009; Vicoso and Bachtrog 2011 for ineffective DC in some organisms). In *Drosophila*, the so-called male-specific lethal (MSL) binds to the chromatin entry sites (CES) and triggers lysine acetylation at the 16th residue in histone H4 (H4K16ac), which in turn stimulates entire male-X upregulation (i.e., global DC) (Alekseyenko et al. 2012; McElroy et al. 2014; Valsecchi et al. 2021). This global DC seems to be also present in the young sex chromosomes, neo-sex chromosomes (hereafter, neo-X and neo-Y, respectively), in *D. miranda* (Fig. 1), as well as in *D. melanica* and *D. robusta*, by co-opting transposable elements into CES (Ellison and Bachtrog 2013; Zhou et al. 2013; Ellison and Bachtrog 2019).

**Fig. 1:**
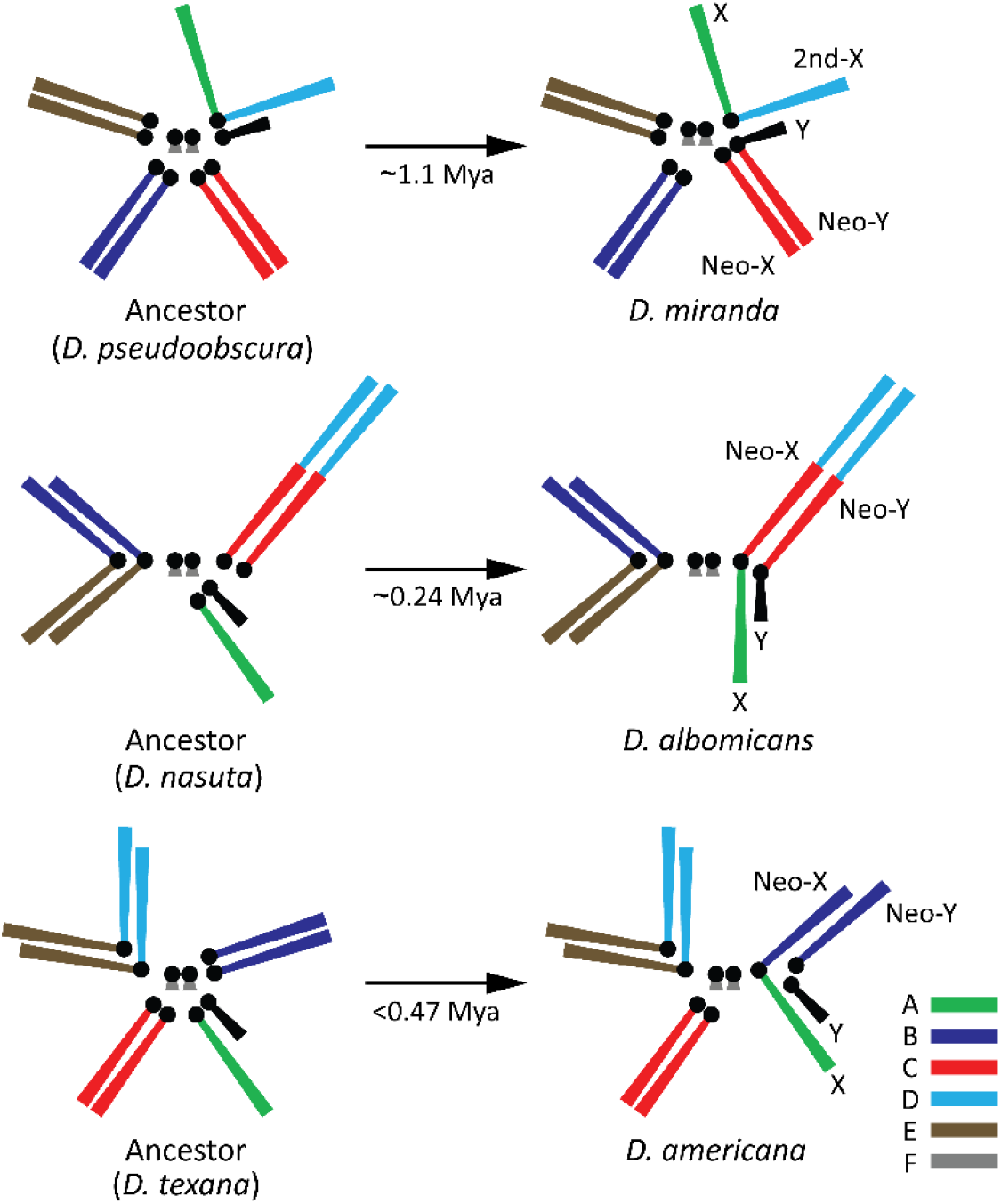
Karyotypes of the three *Drosophila* species and their ancestors. Male karyotypes are shown. Each chromosome color corresponds to the color of Muller elements shown on the bottom-right corner.

In addition to global DC, Nozawa et al. (2018) reported that individual neo-X-linked genes may be upregulated in response to pseudogenization of neo-Y-linked homologs in *D. miranda* [i.e., gene-by-gene or localized DC; see also Nozawa et al. (2014)]. The presence of gene-by-gene DC seems particularly important at the initial stage of sex chromosome evolution when global DC has not yet been established, to counteract the deleterious effects caused by the sequential losses of individual neo-Y-linked genes. However, the pseudogenization rate is accelerated on not only neo-Y, but also neo-X in *D. miranda* (Nozawa et al. 2016), implying that acquiring sex chromosomes may potentially be more deleterious than previously recognized. Yet, whether the observations in *D. miranda* are true for other species and there is a common evolutionary trajectory of sex chromosomes remains unexplored. To address this question, we need to investigate additional young sex chromosomes with independent origins.

In this study, we therefore investigated two other neo-sex chromosomes that originated independently in the *Drosophila* lineages. One such neo-sex chromosome emerged in *D. albomicans* ∼0.24 million years ago (Mya) after the divergence from *D. nasuta* (Satomura and Tamura 2016; Wei and Bachtrog 2019; Mai et al. 2020). The neo-sex chromosome in this species emerged via the fusion of each chromosome 3 (Muller elements C and D) to the canonical X and Y, respectively (Fig. 1). The other neo-sex chromosome focused on in this study is present in *D. americana*. The neo-sex chromosome in this species is likely to have emerged <0.47 Mya after splitting from its sibling species, *D. texana* (Vieira et al. 2003). One of the chromosome 4 (Muller element B) was fused to X and became neo-X, and consequently, the other unfused chromosome 4 became neo-Y (Fig. 1). We also re-investigated the neo-sex chromosome (Muller element C) in *D. miranda*, because its genome sequence has greatly been updated (Mahajan et al. 2018; Bachtrog et al. 2019) after the previous studies (Nozawa et al. 2016; 2018).

Analyzing the neo-sex chromosomes in the three species and the orthologous autosomes in their closely-related species, we here report the evolutionary trajectories of the three independent neo-sex chromosomes. In particular, we focused on whether gene-by-gene DC and accelerated pseudogenization on neo-X are shared among the three *Drosophila* species. It should be mentioned that Muller element C became the neo-sex chromosomes independently in *D. miranda* and *D. albomicans* (Fig. 1). Since genes on the same Muller element are largely orthologous, we also addressed whether parallel evolution occurred at the gene level in these two neo-sex chromosomes.

## RESULTS

### Genomes and transcriptomes of the nine *Drosophila* species

Using long-read sequencers (PacBio RS II and Sequel; Pacific Biosciences, Menlo Park) in combination with a short-read sequencer (HiSeq 2500; Illumina, San Diego), we sequenced and assembled the genomes of *D. albomicans, D. nasuta, D. kohkoa, D. americana, D. texana*, and *D*.*novamexicana* (Table S1). The PacBio RS II-based genome sequence of *D. albomicans* (strain no. 15112-1751.03 from Nankung, Taiwan) has already been published (Mai et al. 2020), but we *de novo* sequenced the genome of another strain (NG3 strain from Okinawa, Japan). To obtain the neo-Y assembly of *D. albomicans* and *D. americana*, male and female short-reads were separately mapped onto the genome assembly based on female DNA, and the neo-X sequence was replaced with male-specific single nucleotide polymorphisms (SNPs) and indels to obtain the neo-Y sequence (see *SUPPLEMENTARY METHODS* for details).

For *D. albomicans* and *D. nasuta*, the contig N50 is ∼22.0 Mb and ∼17.9 Mb, respectively (Table S2), roughly equivalent to the average length of euchromatic regions of a Muller element (Celniker et al. 2002). For other species, the contig N50 is ∼1.6 to ∼3.6 Mb, shorter than the N50 of the former two species. However, the number of contigs was less than 1,000 in all sequenced genomes (as small as 87 in *D. kohkoa*). To evaluate these genome assemblies, we used benchmarking universal single-copy orthologs (BUSCO) ver. 4.0.4 (Seppey et al. 2019). For all nine genome assemblies including *D. miranda, D. pseudoobscura*, and *D. obscura*, more than 98% of the 3,285 single-copy genes in dipteran species were identified as complete open reading frames (ORFs) (Table S3).

To determine the transcriptomes, we conducted the RNA sequencing (RNA-seq) from whole bodies of females and males at the larval, pupal, and adult stages (Table S4). The number of functional protein-coding genes ranged from ∼9,700 to ∼13,900 depending on the species (Table S5). To estimate the expression level of each gene, we also conducted RNA-seq in the ovary and testis (Table S4). The chromosomal location, expression level, and functionality of each gene and transcript are listed in Tables S6 and S7, respectively.

### Proportion of pseudogenes on neo-sex chromosomes

Bachtrog et al. (2019) reported that the number of genes on Y/neo-Y significantly increased after becoming neo-sex chromosomes in *D. miranda*. We also confirmed that the number of genes on Y/neo-Y was ∼3.4 times greater than that on neo-X (Fig. 2A). However, ∼80% of the genes were classified as pseudogenes due to ORF disruption and/or cessation of transcription. The number of functional genes was slightly smaller on neo-Y than that on neo-X (1,862 and 2,082, respectively). When all inparalogs for each category (i.e., functional, silenced, disrupted, silenced-disrupted, and unclassified genes) were put together, the number of functional gene groups on neo-Y became much smaller than that on neo-X (1,439 and 2,031 groups, respectively; Fig. S1). Therefore, although the total number of genes on Y/neo-Y is significantly greater than that on neo-X, Y/neo-Y has considerably degenerated in *D. miranda* and the functional repertoire on Y/neo-Y is smaller than that on neo-X.

**Fig. 2:**
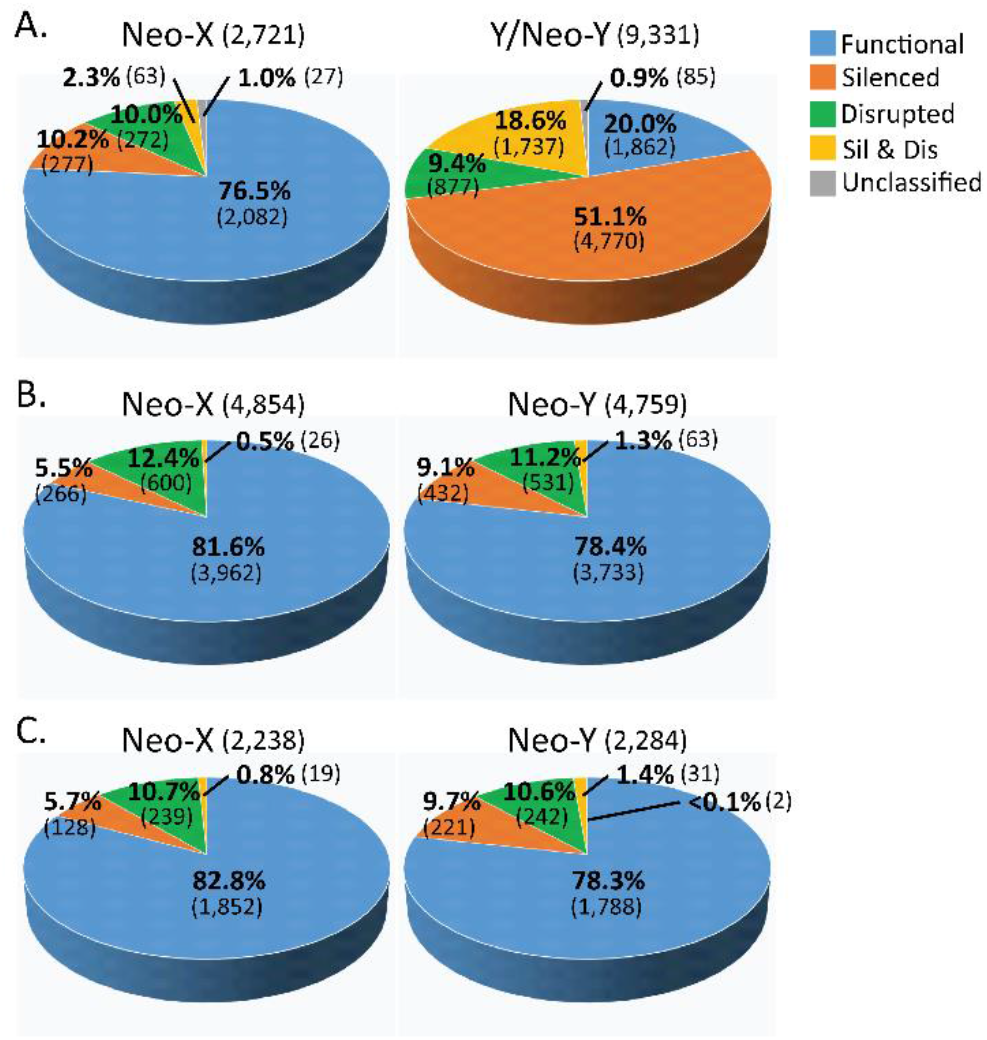
Classification of neo-X- and neo-Y-linked genes in (A) *Drosophila miranda*, (B) *D. albomicans*, and (C) *D. americana*. The numbers in parentheses indicate the numbers of genes in each category. Colors on each pie chart correspond to the categories shown on the top-right corner. Silenced, disrupted, and silenced-and-disrupted genes are pseudogenes.

In contrast, the gene classifications of neo-X and neo-Y were similar in *D. albomicans* and *D. americana*, reflecting their recent origins compared with *D. miranda* (Fig. 2B-C). In addition, the number of orthologous gene groups was similar to the number of genes (Fig. S1), indicating that gene duplication was not common on the neo-sex chromosomes after their emergence in these two species. Therefore, degeneration and specialization of neo-sex chromosomes have less proceeded in these two species compared with those in *D. miranda* (see also Wei and Bachtrog 2019).

### Gene-by-gene and global DC on neo-Xs

Using the *R*_Lin_ index (Lin et al. 2012), Nozawa et al. (2018) reported gene-by-gene DC on the *D. miranda* neo-X. Specifically, the DC level (i.e., the *R*_Lin_ value) for the neo-X-linked genes with pseudogenized neo-Y-linked homologs (X_F_-Y_P_) was significantly greater than that for the neo-X-linked genes with functional neo-Y-linked homologs (X_F_-Y_F_). The updated datasets confirmed this pattern in all tissues examined in *D. miranda* (Fig. 3A). In *D. albomicans*, the trend was the same as that in *D. miranda*, but the difference in the *R*_Lin_ value between the X_F_-Y_F_ and X_F_-Y_P_ groups was significant only in the larva and adult, but not in the pupa and testis (Fig. 3B). In *D. americana*, the difference was significant in the pupa and adult, but not in others (Fig. 3C). Therefore, the gene-by-gene DC seems to be also present on neo-Xs in the two species, but less conspicuous compared with *D. miranda*.

**Fig. 3:**
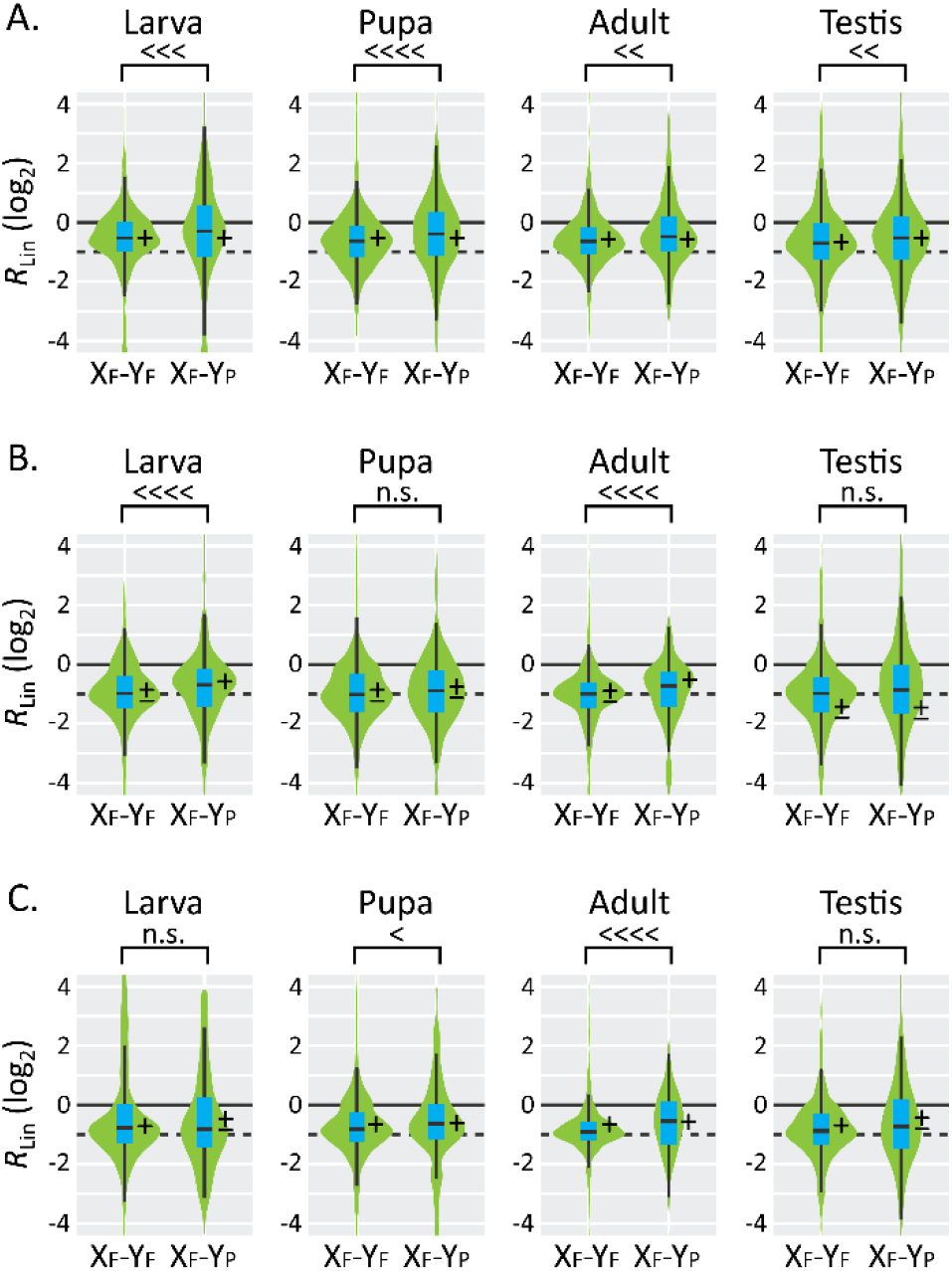
Relationship between the functionality of neo-Y-linked genes and the DC of neo-X-linked homologs in the larva, pupa, adult, and testis (from left to right) of (A) *Drosophila miranda*, (B) *D. albomicans*, and (C) *D. americana*. For computing *R*_Lin_, the corrected fragments per kilobase of exon per million mapped reads (cFPKM, FPKM normalized by median, see *SUPPLEMENTARY METHODS* for details) was used. X_F_-Y_F_, a group of genes with functional neo-X-linked and neo-Y-linked homologs; X_F_-Y_P_, a group of genes with functional neo-X-linked homologs and but pseudogenized neo-Y-linked homologs. The numbers of genes analyzed in the X_F_-Y_F_ and X_F_-Y_P_ groups are 664 and 435 for *D. miranda*, 2,449 and 218 for *D. albomicans*, and 1,204 and 93 for *D. americana*, respectively. A box plot is also shown on each violin plot. Differences in median between groups were tested based on a permutation test with 10,000 replicates under the null hypothesis of *R*_Lin_ (X_F_-Y_F_) = *R*_Lin_ (X_F_-Y_P_): <<<<, *P*<10^−4^; <<<, *P*<10^−3^; <<, *P*<0.01; <, *P*<0.05; n.s., *P*≥0.05. A solid line indicates the *R*_Lin_ value of 1 (0 in log_2_) indicating perfect DC, whereas a broken line corresponds to a value of 0.5 (−1 in log_2_) indicating no DC. +, –, and ± along each plot means that the median *R*_Lin_ value is >0.5, <0.5, and 0.5 at the 5% significance level, respectively, based on a bootstrap test with 10,000 replications.

Another difference among the neo-Xs in these species is the *R*_Lin_ value for the X_F_-Y_F_ group. As already mentioned, the MSL recognizes the CES, which triggers histone acetylation to initiate global DC on the male neo-X in *D. miranda* (Ellison and Bachtrog 2013). Therefore, in *D. miranda*, not only X_F_-Y_P_ genes, but also X_F_-Y_F_ genes showed *R*_Lin_ values significantly greater than 0.5, the expected *R*_Lin_ value under no DC (Fig. 3A) (see also Nozawa et al. 2018). In contrast, X_F_-Y_F_ gene upregulation was not necessarily conspicuous in *D. albomicans* and *D. americana*. In particular, *R*_Lin_ for the X_F_-Y_F_ genes was ∼0.5 in all samples examined in *D. albomicans* (Fig. 3B). Therefore, global DC is unlikely to be established in *D. albomicans* or *D. americana*. It should be mentioned that when corrected transcripts per kilobase million mapped reads (cTPM, TPM normalized by the median TPM for each sample; Fig. S2) was used to compute *R*_Lin_ instead of cFPKM (Fig. 3) for estimating gene expression levels, the results were qualitatively same.

### Accelerated pseudogenization on neo-Xs

Next, we examined whether the accelerated pseudogenization on neo-X as well as neo-Y reported in *D. miranda* (Nozawa et al. 2016) occurred on the neo-sex chromosomes in *D. albomicans* and *D. americana*. First, we confirmed the accelerated pseudogenization on both neo-X and neo-Y in *D. miranda* with the updated datasets (Fig. 4A–B). The pseudogenization rate on the neo-X lineage (branch *a*) was significantly greater than that on the orthologous autosome in *D. pseudoobscura* (branch *e*) (*P*=1.5×10^−6^ by χ^2^ test). Similarly, the pseudogenization rate on the neo-X lineage (branch *a*) was significantly greater than that on the *D. miranda* ancestral lineage before the emergence of neo-sex chromosome (branch *c*) (*P*=5.5×10^−10^). The sum of pseudogenization events on branches *a* and *c* was also significantly greater than the sum of those on branches *d* and *e* (*P*=2.9×10^−3^).

**Fig. 4:**
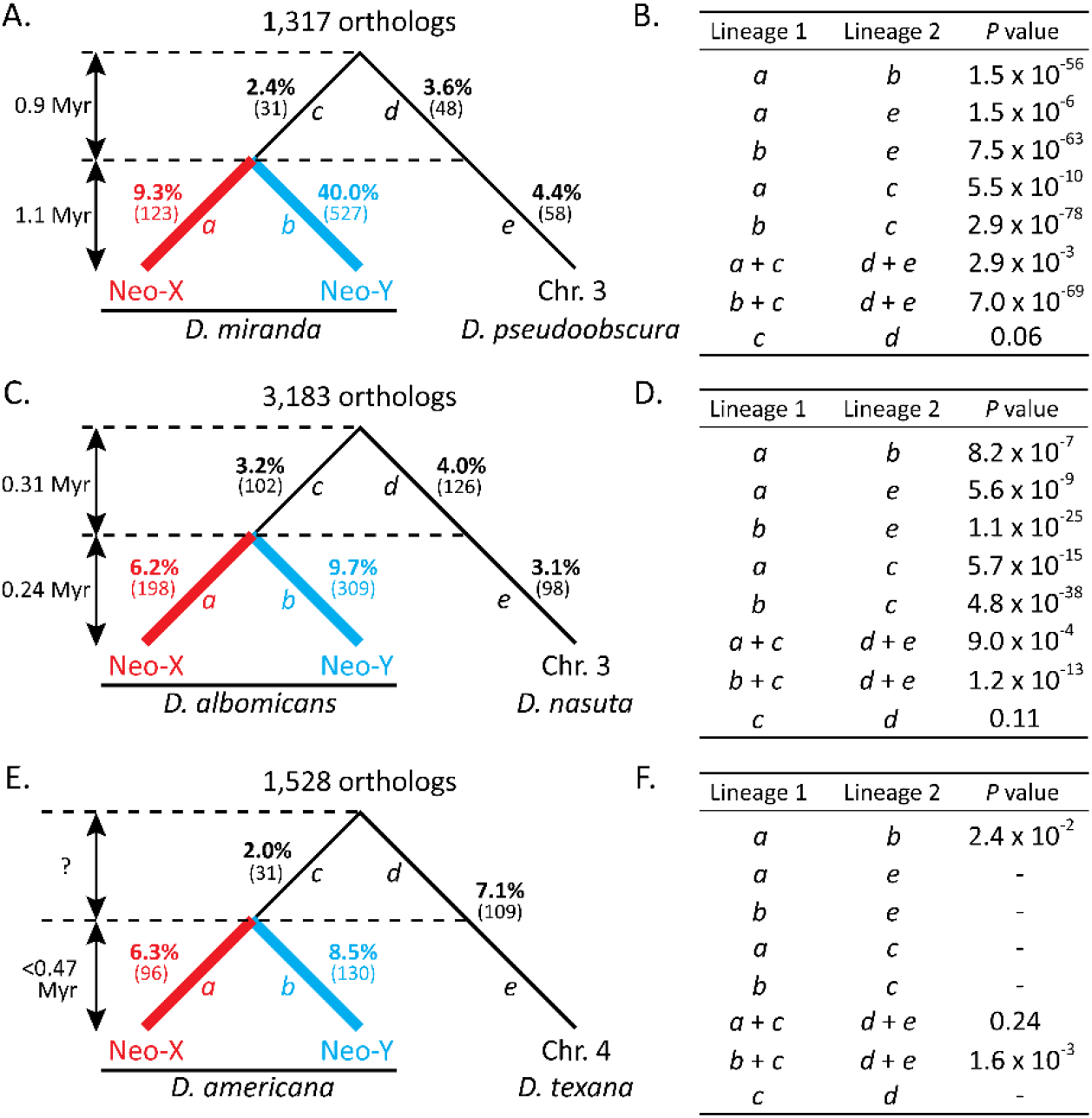
Pseudogenization events before and after neo-sex chromosome emergence. (A, C, and E) Proportions of genes that were pseudogenized in (A) *Drosophila miranda*, (C) *D. albomicans*, and (E) *D. americana* lineages compared with those in their closely-related species. The numbers in parentheses indicate the number of genes pseudogenized in each lineage. The orthologs regarded as functional in the outgroup species (*D. obscura, D. kohkoa*, and *D. novamexicana*, respectively) and located on the same Muller element without any inparalogs were used for this analysis. The number of such orthologs is shown above each tree. (B, D, and E) Statistical significance in the difference of the pseudogenization rate between lineages of (B) *D. miranda*, (D) *D. albomicans*, and (F) *D. americana* by χ^2^ test under the null hypothesis of equal pseudogenization rates between lineages. Lineages *a* to *e* correspond to the branches in A, C, and E.

Similar patterns were also observed in the *D. albomicans* lineage (Fig. 4C–D). Differences in the pseudogenization rates on the branches *a* and *e*, the branches *a* and *c*, and branches *a* + *c* and *d* + *e* were all statistically significant. In contrast, the difference was not significant in the *D. americana* lineage (96 + 31 = 127 for *a* + *c* and 109 for *d* + *e, P*=0.24), although the trend was qualitatively same (Fig. 4E–F). The difference in the pseudogenization rates on neo-X and neo-Y was also less clear (*P*=0.024), probably reflecting their recent origin. It should be noted that the pseudogenization rate was not significantly different for most of other chromosomes between the species with neo-sex chromosomes and their close relatives without neo-sex chromosomes (Fig. S3). Therefore, the mutation rate as well as general functional constraints between the former and the latter species would be essentially the same. These results indicate that the three species with neo-sex chromosomes share an evolutionary trajectory with respect to the accelerated pseudogenization on neo-X, although the extent varies with species, possibly due to the age difference among the neo-sex chromosomes.

### Pseudogenization pattern on neo-sex chromosomes

We next examined pseudogenization pattern on neo-X and neo-Y. First, to test whether pseudogenes on the neo-sex chromosomes were under less functional constraints before being linked to neo-sex chromosomes, we examined the relationship between functionality of neo-sex-linked genes and ratio of nonsynonymous to synonymous nucleotide divergence per site (*d*_N_/*d*_S_) (Fig. 5). In the *D. miranda* lineage, pseudogenes on neo-X (i.e., X_P_-Y_F_ and X_P_-Y_P_ genes) tended to have less functional constraints when they were functional in the ancestral lineage before the emergence of neo-X and neo-Y (branch *c* in Fig. 5A). A similar but less clear tendency was also observed in the *D. albomicans* and *D. americana* lineages (branch *c* in Fig. 5B–C). X_P_-Y_F_ genes tend to have evolved faster than X_F_-Y_F_ and X_F_-Y_P_ on the neo-X lineages in all three species as expected (branch *a* in Fig. 5A-C). However, X_P_-Y_F_ genes tend to have evolved even faster than X_P_-Y_P_ genes, particularly in the *D. miranda* lineage (middle panel in Fig. 5A). X_F_-Y_P_ genes, in contrast, did not necessarily evolve faster than other gene categories in the ancestral lineage (branch *c*), although they have evolved faster on the neo-Y lineages (branch *b*) as expected (Fig. 5A–C). Therefore, neo-X-specific pseudogenization events have primarily occurred because these genes were under less functional constraints before becoming neo-sex-linked, whereas neo-Y-specific pseudogenization events are unlikely to be associated with functional constraints in the ancestor very much.

**Fig. 5:**
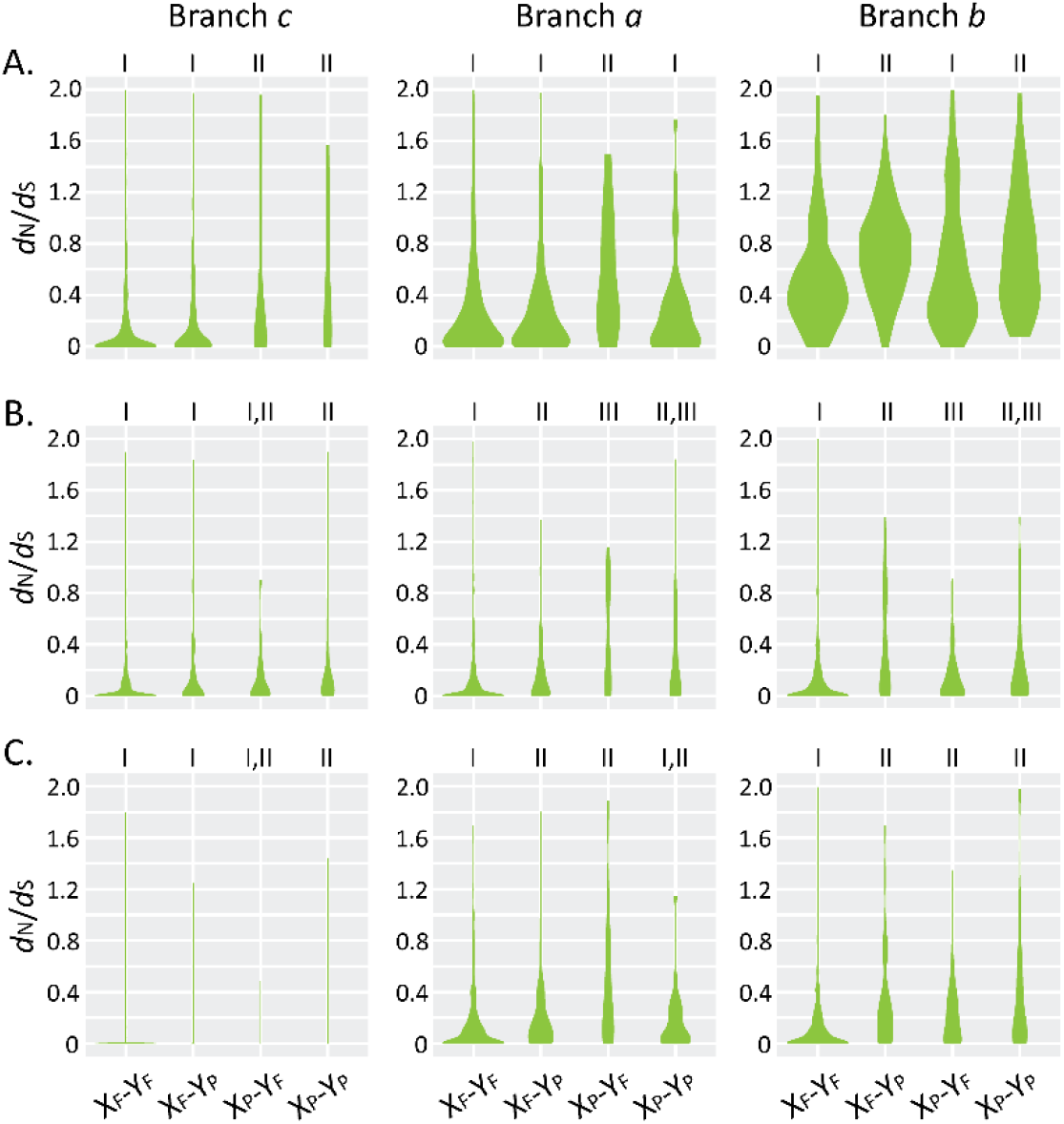
Relationship between functionality of neo-sex-linked genes and ratio of nonsynonymous to synonymous nucleotide divergence per site (*d*_N_/*d*_S_) in (A) *Drosophila miranda*, (B) *D. albomicans*, and (C) *D. americana* lineages. Branches *c* (left), *a* (center), and *b* (right) corresponds to those in Fig. 4. Violin plots were used to show *d*_N_/*d*_S_ distribution. The orthologs regarded to be functional in the outgroup species (*D. obscura, D. kohkoa*, and *D. novamexicana*, respectively) and located on the same Muller element without any inparalogs were used for this analysis. The number of genes analyzed for each category is as follows: 681 genes for X_F_-Y_F_, 474 genes for X_F_-Y_P_, 70 genes for X_P_-Y_F_, and 92 genes for X_P_-Y_P_ in the *D. miranda* lineage; 2,551 genes for X_F_-Y_F_, 277 genes for X_F_-Y_P_, 166 genes for X_P_-Y_F_, and 189 genes for X_P_-Y_P_ in the *D. albomicans* lineage; 1,267 genes for X_F_-Y_F_, 119 genes for X_F_-Y_P_, 85 genes for X_P_-Y_F_, and 57 genes for X_P_-Y_P_ in the *D. americana* lineage. Differences in median values between categories were tested based on a permutation test with 10,000 replicates under the null hypothesis of the equal *d*_N_/*d*_S_ ratio between categories. Same roman numerals indicate *P*≥0.05, whereas different numerals mean *P*<0.05 between categories.

Next, we conducted gene ontology (GO) analysis to extract features in functional genes or pseudogenes on neo-sex chromosomes. The analysis detected many GO terms enriched in functional genes and pseudogenes on neo-X and neo-Y (Tables S8–19). However, the trend was largely shared between neo-X and neo-Y of the three species. For example, genes involved in metabolic processes tended to be functional on both neo-X and neo-Y of the three species. On the contrary, genes related to detection of chemical stimuli commonly tended to be nonfunctional on the three neo-sex chromosomes. The shared feature between neo-X and neo-Y in each species may be expected, because they had shared evolutionary history until recently. However, this explanation would be insufficient for the enriched GO terms shared among species. We speculate that the above patterns seem to be general features applicable to all chromosomes. Indeed, chemosensory receptor genes involved in detecting chemical stimuli are known to have been under dynamic turn-over and pseudogenization in many animals (Niimura and Nei 2007; Nozawa and Nei 2007; Nei et al. 2008; Hayakawa et al. 2014).

Therefore, we next examined whether pseudogenization events are associated with spatiotemporal gene expression patterns. For this analysis, genes were classified based on the tissue where the genes were expressed the highest among the samples examined. Ideally, the gene expression level in the ancestral species without neo-sex chromosomes should be used to classify the genes, but obtaining RNA from ancestral species is impossible. We therefore used the gene expression level in the closely-related species without neo-sex chromosomes as a proxy for the gene expression level in the ancestor (i.e., *D. pseudoobscura, D. nasuta*, and *D. texana* for *D. miranda, D. albomicans*, and *D. americana*, respectively).

On neo-X, the proportion of pseudogene was not significantly different among groups in all three lineages (second panel from the right in Fig. 6A–C). On canonical X, the genes expressed the highest in the ovary showed a consistent trend of maintaining their functions, although the trend was statistically significant only in *D. miranda* (second panel from the left in Fig. 6A–C). A similar but non-significant pattern was also observed in the *D. miranda* and *D. americana* neo-Xs [Fig. 6A, C, see also Nozawa et al. (2016)]. Therefore, neo-X may be in a transitional state toward canonical X with respect to gene content.

**Fig. 6:**
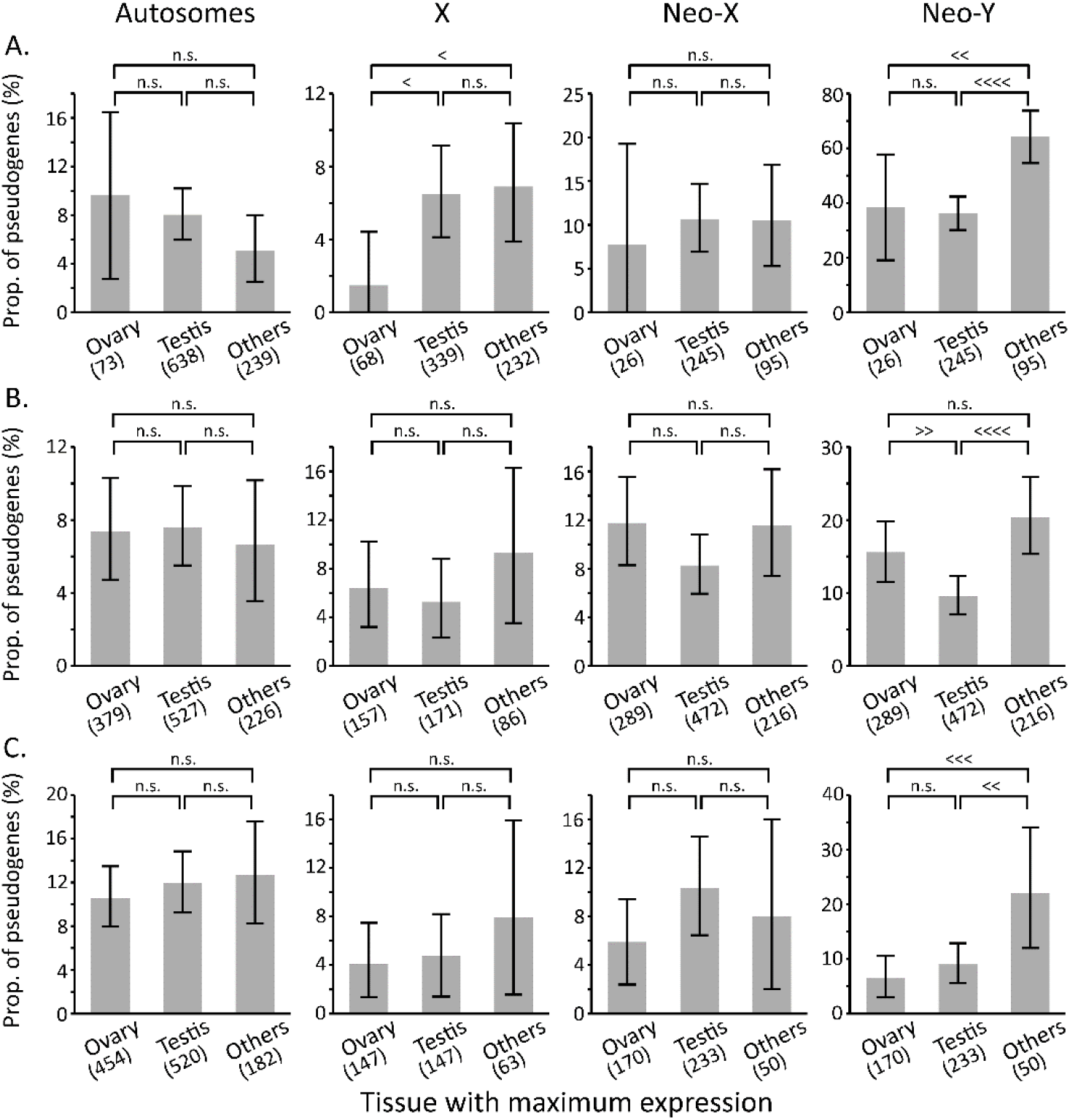
Relationship between spatiotemporal gene expression pattern and pseudogenization. All genes were classified into three groups based on the tissue with the highest expression in (A) *Drosophila pseudoobscura*, (B) *D. nasuta*, and (C) *D. texana*, the closely-related species of *D. miranda, D. albomicans*, and *D. americana*, respectively, based on the cFPKM value. However, to remove the genes with similar cFPKMs in multiple tissues, only genes with at least two-fold cFPKM in a tissue compared with that in the other tissues examined were used for this analysis. The orthologs regarded to be functional in the close relatives (*D. pseudoobscura* and *D. obscura* for *D. miranda, D. nasuta* and *D. kohkoa* for *D. albomicans*, and *D. texana* and *D. novamexicana* for *D. americana*, respectively) and located on the same Muller element without any inparalogs were used for this analysis. The number of genes in each category is shown in parenthesis. Error bars show the 95% confidence interval based on a bootstrap resampling with 10,000 replicates. Statistical significance between groups was calculated by a permutation test with 10,000 replicates: <<<< or >>>>, *P*<10^−4^; <<< or >>>, *P*<10^−3^; << or >>; *P*<0.01; < or >, *P*<0.05; n.s., *P*≥0.05.

On neo-Y, the genes expressed the highest in the testis showed a significantly lower proportion of pseudogenes than those expressed the highest in somatic tissues for all three lineages (rightmost panel in Fig. 6A–C), consistent with previous studies in *D. miranda* (Kaiser et al. 2011; Nozawa et al. 2018). The genes expressed the highest in the ovary also tended to maintain their functions even after being neo-sex-linked in all three lineages, which was also consistent with the finding in *D. miranda* (Nozawa et al. 2018). It should be mentioned that there was no clear-cut difference among tissues on autosomes (leftmost panel in Fig. 6A–C), probably because of biparental inheritance with equal contribution from males and females. Notably, the general pattern remained unchanged even when TPM rather than cFPKM was used to estimate the gene expression level (Fig. S4). Therefore, pseudogenization pattern shared on neo-Y in the three species would be primarily due to the linkage of the genes to Y.

### Parallel pseudogenization on the neo-sex chromosomes in *D. miranda* and *D. albomicans*

The above analyses consistently indicate that the three neo-sex chromosomes largely share a common evolutionary trajectory after their independent emergence. Since Muller element C independently became the neo-sex chromosomes in *D. miranda* and *D. albomicans*, we directly examined whether the same orthologs tend to have been parallelly pseudogenized in both lineages (Fig. 7). The result showed that three orthologs were parallelly pseudogenized on the neo-X lineages, which was within a range under random pseudogenization on each lineage (Fig. 7A and D). In contrast, there were 35 parallel pseudogenization events on the neo-Y lineages, which was significantly greater than the distribution under random pseudogenization (Fig. 7B and E). It should be noted that the same genes tend to have also been pseudogenized on Muller elements A, D, and E, but the trend was much weaker compared with the pattern on the neo-Y lineages of the two species (Fig. S5). It should also be noted that there was no parallel pseudogenization on the Muller element C in the ancestral lineage before the emergence of the neo-sex chromosomes (Fig. 7C and F). Therefore, parallel pseudogenization on the two neo-Ys is unlikely to be accidental but likely reflects a common evolutionary trajectory regarding pseudogenization pattern on neo-Y.

**Fig. 7:**
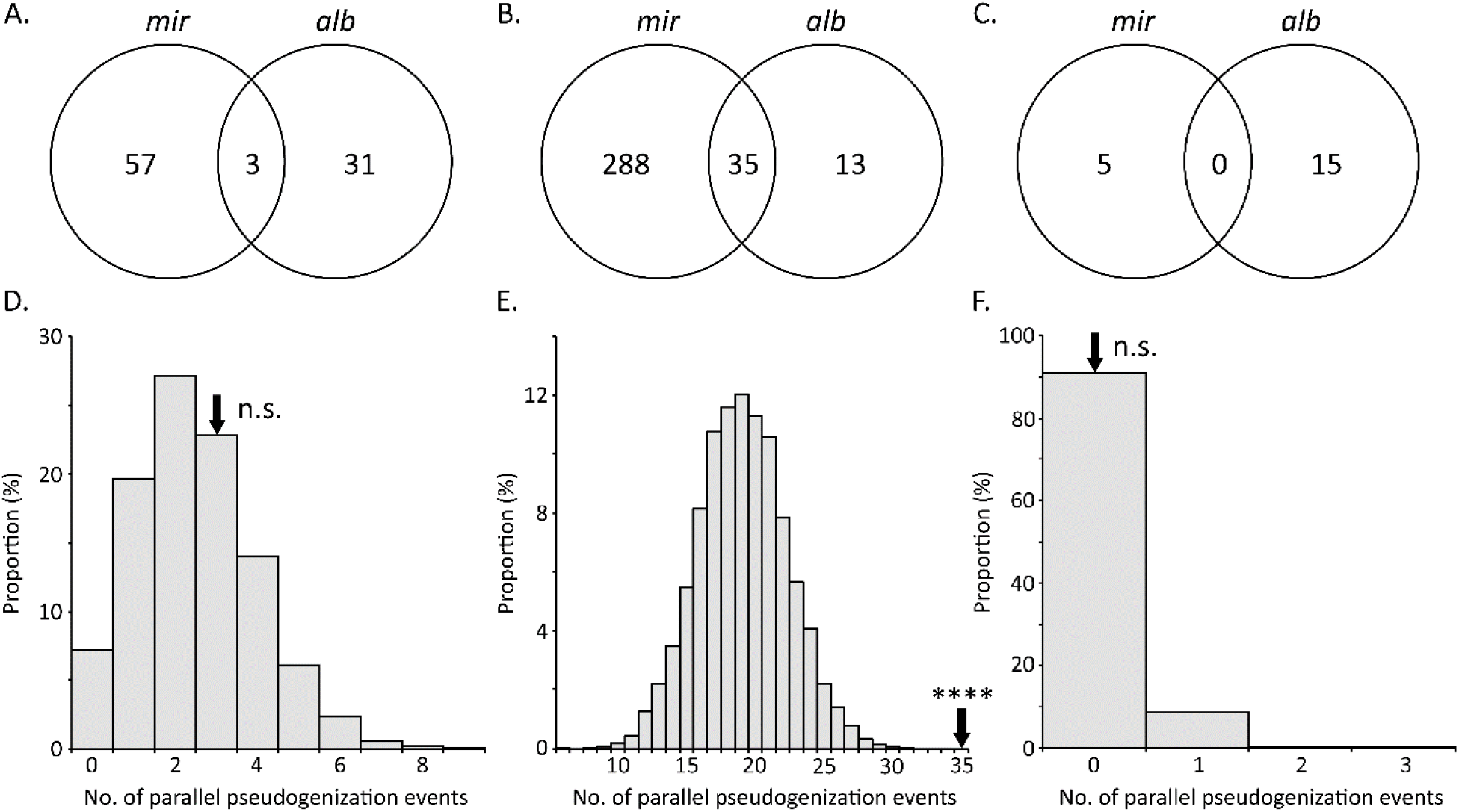
Parallel pseudogenization events on the neo-sex chromosomes in *Drosophila miranda* (*mir*) and *D. albomicans* (*alb*) lineages. A total of 814 orthologs regarded to be functional in the outgroup species (*D. obscura* and *D. kohkoa*, respectively) and located on the Muller element C without any inparalogs were used for the analysis. A Venn diagram shows the number of lineage-specific or parallel pseudogenization events on the (A) neo-X, (B) neo-Y, and (C) ancestral lineages (i.e., branches *a, b*, and *c*, respectively, in Figs. 4A and 4C) of *D. miranda* and *D. albomicans*. Distribution of the number of pseudogenization events based on the permutation test with 10,000 replicates under the assumption of random pseudogenization on the (D) neo-X, (E) neo-Y, and (F) ancestral lineages (i.e., branches *a, b*, and *c*, respectively, in Figs. 4A and 4C) of *D. miranda* and *D. albomicans* is shown below each Venn diagram. An arrow represents the observed number of parallel pseudogenization events with the following statistical significance: ****, *P*<10^−4^; n.s., *P*≥0.05.

We further searched for unique features of the genes that experienced parallel pseudogenization. The *d*_N_/*d*_S_ ratios for the genes that experienced parallel pseudogenization and lineage-specific pseudogenes were similar (Fig. S6), although the ratios for the pseudogenes tended to be higher than those for functional genes on the neo-Y lineages (branch *b* in Fig. S6). However, the orthologs of the neo-Y pseudogenes in the closely-related species (*D. pseudoobscura* for *D. miranda* and *D. nasuta* for *D. albomicans*, respectively) showed tissue-specific expression as indicated by *τ* (Yanai et al. 2005) at a significantly greater extent than those of functional genes, although there was no significant difference in *τ* between orthologs of lineage-specific and shared pseudogenes (Fig. 8A). The number of expressed tissues also tended to be smaller in the orthologs of shared pseudogenes than those of functional genes and lineage-specific pseudogenes on neo-Y (Fig. 8B). Moreover, the orthologs of shared neo-Y-linked pseudogenes in *D. pseudoobscura*, closely related to *D. miranda*, tended to show female-biased expression compared with the orthologs of functional genes and lineage-specific pseudogenes at the larval and pupal stages in *D. miranda* (Fig. 8C), although the pattern was less clear in the *D. albomicans* lineage (Fig. 8D). Therefore, the female-biased genes expressed only in a small number of tissues tend to be parallelly pseudogenized on neo-Y.

**Fig. 8:**
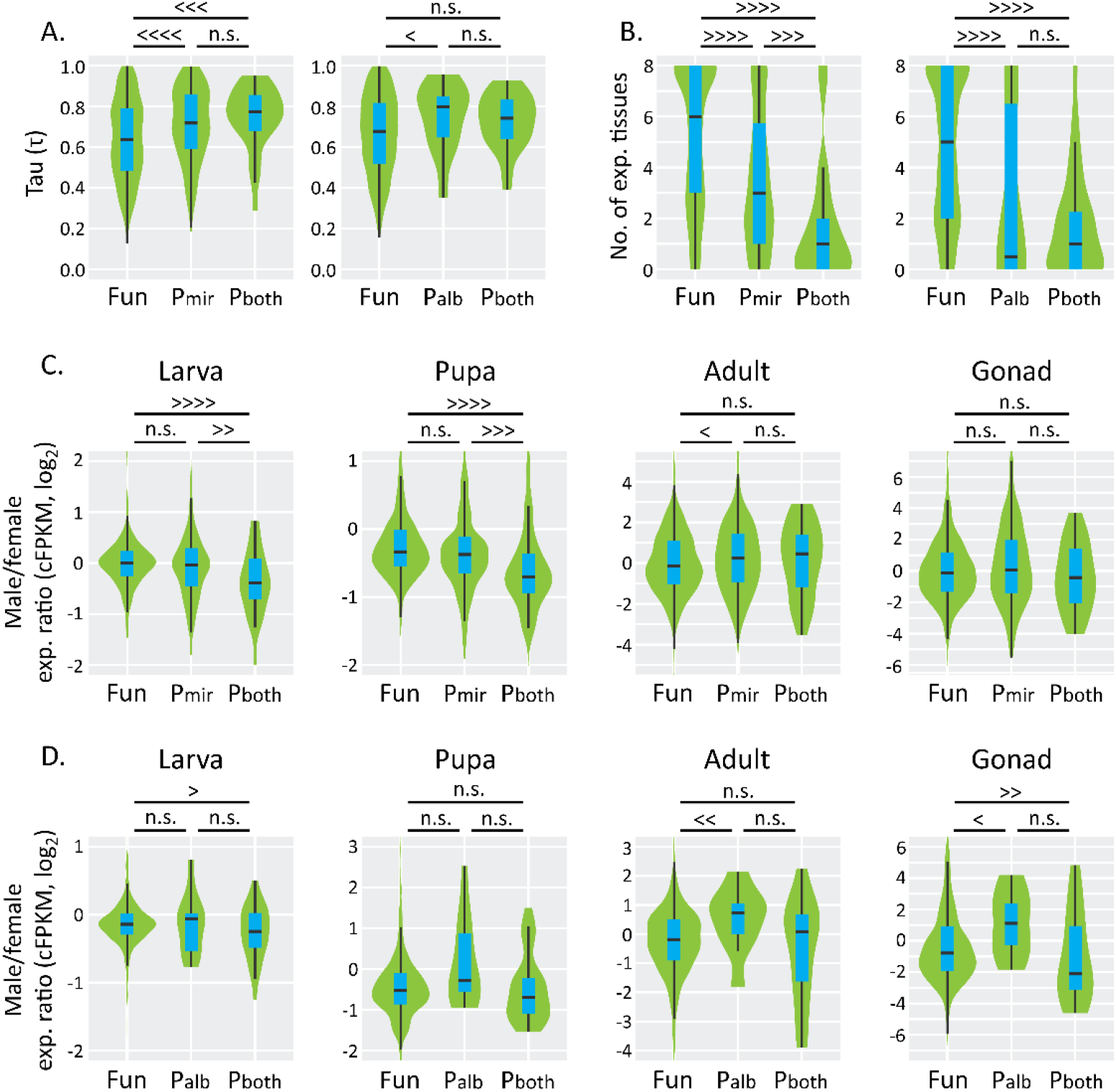
Relationship between parallel pseudogenization on neo-Y and their expression pattern. (A) Relationship between pseudogenization pattern and tissue specificity in gene expression (*τ*). *τ* was computed for orthologs in *D. pseudoobscura* (left panel) and *D. nasuta* (right panel), the closely-related species of *D. miranda* and *D. albomicans*, respectively, as proxies of ancestral species. (B) Relationship between pseudogenization pattern and the number of expressed tissues for each gene. A gene was regarded to be expressed in a tissue if the cFPKM was ≥1 in *D. pseudoobscura* (left panel) and *D. nasuta* (right panel). (C, D) Relationship between pseudogenization pattern and male to female gene expression ratio based on the cFPKM value. Fun, neo-Y-linked genes functional at least in *D. miranda* (left panel in A and B and all panels in C) or *D. albomicans* (right panel in A and B and all panels in D); P_mir_, neo-Y-linked genes nonfunctional in *D. miranda* but functional in *D. albomicans*; P_alb_, neo-Y-linked genes nonfunctional in *D. albomicans* but functional in *D. miranda*; P_both_, neo-Y-linked genes nonfunctional in both *D. miranda* and *D. albomicans*. Differences in median between categories were statistically evaluated based on a permutation test with 10,000 replicates: <<<< or >>>>, *P*<10^−4^; <<< or >>>, *P*<10^−3^; << or >>; *P*<0.01; < or >, *P*<0.05; n.s., *P*≥0.05.

## DISCUSSION

A variety of sex-determination systems have come into existence in a long history of life. Among them, sex chromosomes have particularly been widespread and repeatedly emerged in many lineages. However, if many gene losses from Y are considered, sex chromosome acquisition is not a simple evolutionary process. In this study, we investigated if there is a common evolutionary trajectory to avoid the neo-sex chromosomes being evolutionary dead-end, using the three *Drosophila* species that independently acquired neo-sex chromosomes and their closely-related species without neo-sex chromosomes. The shared features that we found in this study can be summarized as follows: 1) The gene-by-gene or localized DC operates on neo-X-linked genes. 2) The pseudogenization rate was accelerated on neo-X. 3) The neo-Y-linked genes with the highest expression in the testis or ovary tend to remain functional. 4) The same orthologs tend to be parallelly pseudogenized on neo-Y in *D. miranda* and *D. albomicans*.

### DC on neo-X in *Drosophila*

When individual Y-linked genes are pseudogenized, immediate upregulation of X-linked homologs would be critical to counteract the deleterious effects of the pseudogenization. In this study, we found the gene-by-gene upregulation of neo-X-linked homologs in the three species with independent origins of neo-sex chromosomes, which has likely played an important role in diminishing the effect of losing neo-Y genes. However, the gene-by-gene DC was less clear in *D. albomicans* and *D. americana*, with younger neo-sex chromosomes, compared with the DC in *D. miranda* with older neo-sex chromosomes. These results imply that the time to fully develop the gene-by-gene DC mechanism is insufficient for the former two species. It should be mentioned that this implication is not necessarily consistent with the previous report that a certain portion of the neo-X-linked genes already have a potential of upregulation (i.e., gene-by-gene DC) before losing their neo-Y homologs in *D. miranda* (Nozawa et al. 2018). Yet, their analysis is unlikely to be conclusive as they also noticed large sampling errors in estimating the pseudogenization time for each neo-Y-linked gene due to the small sequence divergence between the neo-sex chromosomes in *D. miranda*. Hence, we did not conduct the same analysis for *D. albomicans* and *D. americana*, in which the sequence divergence between neo-X and neo-Y is even smaller. Therefore, how quickly the mechanism of gene-by-gene DC is established after neo-Y-linked gene loss remains unclear. We are currently making the mutants of *D. miranda* and *D. albomicans* in which several functional neo-Y-linked genes are knocked out and see if the neo-X-linked homologs are immediately upregulated in one generation.

We also found that global DC as well as gene-by-gene DC operate on the neo-X in *D. miranda* as already reported (Ellison and Bachtrog 2013; Nozawa et al. 2018), whereas in *D. albomicans* and *D. americana*, global DC is unlikely to exist. This observation strengthens the speculation by a previous study (Nozawa et al. 2018) that gene-by-gene DC is developed first in the early stage of sex chromosome evolution and global DC is developed later and gradually takes over the role of gene-by-gene DC as Y degeneration proceeds. Under this scenario, the neo-Xs in *D. albomicans* and *D. americana* are so young that global DC has not been established yet, whereas the neo-X in *D. miranda* is in a transient stage from gene-by-gene to global DC. Consistent with this scenario, the neo-Xs that independently originated in *D. melanica* and *D. robusta* about 4–15 Mya (Flores et al. 2008) have already acquired the MSL binding sites, i.e., the global DC mechanism (Ellison and Bachtrog 2019). On the contrary, the neo-X with an age of ∼0.85 Mya in *D. busckii* is unlikely to acquire the global DC (Zhou and Bachtrog 2015). These observations imply that approximately one million years may be necessary for neo-X to at least partially develop global DC mechanism in *Drosophila* species. It should be mentioned that neo-Y degeneration is not conspicuous in *D. albomicans* and *D. americana*. Therefore, the effects of gene loss from neo-Y on fitness may be limited even without strong gene-by-gene DC.

### Accelerated pseudogenization on neo-X and neo-Y in *Drosophila*

As discussed above, gene-by-gene DC certainly operates on neo-X, which is likely critical at the early stage of sex chromosome evolution for organisms to maintain their fitness. However, gene-by-gene DC is unlikely to completely mitigate the deleterious effects caused by neo-Y-linked gene loss, because the pseudogenization rate is also accelerated on neo-X, at least in *D. miranda* and *D. albomicans* (and the trend is the same in *D. americana* as well). If a gene is pseudogenized or lost from neo-X, gene-by-gene DC cannot operate on its neo-Y-linked homolog. However, the neo-X-linked pseudogenes tend to have been under less functional constraints when they were functional on an autosome. Thus, the fitness effects of these pseudogenizations seem to be minimized.

What kinds of evolutionary forces have driven this accelerated rate of pseudogenization on neo-sex chromosomes? For neo-Y, suppression of meiotic recombination and small effective population size (one-fourths compared with the size of an autosome) would be sufficient explanations. For neo-X, however, the situation is more complex. The effective population size of neo-X is expected to be three-fourths compared with that of an autosome under equal sex ratio. However, unlike many other organisms, most *Drosophila* species do not recombine at male meiosis (John et al. 2016). Therefore, the proportion of chromosomes which recombine at meiosis is two-thirds for neo-X, greater than two-fourths for an autosome (Charlesworth 2012). Therefore, the inclusive efficacy of natural selection on neo-X and an autosome is expected to be similar (i.e., 3/4 × 2/3 = 1/2 for neo-X and 4/4 × 2/4 = 1/2 for an autosome). Therefore, other forces such as sexual conflict may have significant effects on the fates of genes on neo-sex chromosomes. Indeed, genes likely involved in sexual conflict are reported to show a biased pattern of pseudogenization on the neo-X in *D. miranda*, which may have played a certain role in reducing the sexual conflict between males and females due to the acquisition of neo-sex chromosomes (Nozawa et al. 2016; but see also Bachtrog et al. 2019 for neo-sex chromosomes as a new battleground for sexual conflict).

The duration of accelerated pseudogenization on neo-X after neo-sex chromosome emergence would also be an intriguing point to be clarified. In the dipteran lineage, dozens of sex chromosome turn-over have occurred (Vicoso and Bachtrog 2015). Because most Y-linked genes in dipteran species, and probably many other species, are pseudogenized, X, but not Y, will be reversed to an autosome (Meisel 2020). Indeed, a large number of genes remain functional on X in many species (Koerich et al. 2008; Cortez et al. 2014; Zhou et al. 2014; Dupim et al. 2018; Bracewell and Bachtrog 2020). Since chromosome-level assemblies of different ages of X have recently become available in several *Drosophila* species (Mahajan et al. 2018; Bracewell et al. 2019; Wei and Bachtrog 2019; Bracewell and Bachtrog 2020), it may be possible to estimate the transition of the pseudogenization rate of X after its emergence in the future.

### Pseudogenization pattern on neo-sex chromosomes in *Drosophila*

We found that the genes under less functional constraints in the ancestor before neo-sex chromosome emergence are frequently pseudogenized after being neo-X-linked. As already mentioned, this trend would minimize the deleterious effect of acquiring sex chromosomes on the fitness of organisms. In contrast, this trend does not necessarily hold for the neo-Y-linked genes, particularly the genes pseudogenized only on neo-Y. Instead, the spatiotemporal gene expression pattern is likely to be a primary factor in determining the fate of neo-Y-linked genes. Specifically, if a gene is highly expressed in the testis of the closely-related species without neo-sex chromosomes (possibly reflecting the gene expression pattern in the ancestor), it tends to remain functional after being neo-Y-linked. This observation is reasonable because Y is transmitted only through males, so the genes whose functions are specific or biased to males, such as genes highly expressed in the testis, are likely maintained.

In this context, however, our finding that the genes with the highest expression in the ovary have a lower proportion of pseudogenes on neo-Y compared with those with the highest expression in other somatic tissues is counterintuitive, because these genes are quite unlikely to be necessary for males. One possibility is that the genes with the highest expression in the ovary are also important in male tissues. Indeed, the genes with the highest and second highest expression in the ovary and male tissues, respectively, tend to remain functional compared with those with the highest and second highest expression in the ovary and other female tissues, respectively, on the *D. miranda* neo-Y (Nozawa et al. 2018). Although our updated dataset for *D. miranda* with more stringent criteria to select the genes for the analysis did not reproduce such patterns (the proportions of pseudogenes are 0.43 and 0.37 for the former and the latter, respectively, with *P*=0.87 by permutation test with 10,000 replications), we observed a weak but non-significant pattern in *D. albomicans* (0.14 and 0.17, respectively, with *P*=0.40) and a strong tendency in *D. americana* (0.02 and 0.09, respectively, with *P*=0.01). Therefore, some genes with the highest expression in the ovary may also be important for males and remain functional on neo-Y.

The second possibility to be considered is the recombination between neo-X and neo-Y. If neo-X and neo-Y recombine at male meiosis, pseudogenes on neo-Y can theoretically be replaced with neo-X-derived functional genes. Although males do not recombine at meiosis in *D. melanogaster* and many other *Drosophila* species as mentioned above (John et al. 2016), *D. nasuta*, closely related to *D. albomicans*, recombines during male meiosis (Satomura and Tamura 2016). In addition, neo-X and neo-Y in *D. albomicans* are likely to have recombined until recently, supported by a large number of shared polymorphisms between neo-X and neo-Y, although male recombination does not occur at present in *D. albomicans* (Satomura and Tamura 2016; see also Wei and Bachtrog 2019). A large number of shared polymorphisms between neo-sex chromosomes was also reported in *D. miranda* (Nozawa et al. 2018), which might also indicate male recombination in *D. miranda* and/or its recent ancestor. In *D. americana*, available data are insufficient to count the number of shared polymorphisms between neo-X and neo-Y. However, unlike those of *D. miranda* and *D. pseudoobscura*, and *D. albomicans* and *D. nasuta*, the geographical distributions of *D. americana* and the closely-related species *D. texana* partly overlap (Sillero et al. 2014). Therefore, if a hybrid male is backcrossed with a *D. americana* female, a neo-Y pseudogene can be replaced with a functional gene that is highly expressed in the ovary, on the orthologous autosome (i.e., chromosome 4) from *D. texana*.

### Parallel pseudogenization of neo-Y-linked genes in *D. miranda* and *D. albomicans*

This study reports a common evolutionary trajectory of the neo-sex chromosomes in *Drosophila* species. One of the most striking findings among these features may be the parallel pseudogenization of neo-Y-linked genes in *D. miranda* and *D. albomicans*. There is no enriched GO term in these genes with parallel pseudogenization, possibly due to a small number of such genes (only 35 of the 814 genes analyzed). However, a detailed inspection revealed that sexual conflict may be involved in at least some of the parallel pseudogenizations (Table S20). For example, *Idgf6* is known to be involved in egg chamber tube morphogenesis (Zimmerman et al. 2017). In addition, *JhI-26*, which was originally identified as a juvenile hormone-inducible protein, was later reported as a sperm protein known to reduce male fertility under overexpression (Liu et al. 2014). Therefore, these genes may be under sexual conflict and thus may have been pseudogenized after neo-Y-linkage.

In conclusion, neo-sex chromosomes in *Drosophila* with independent origins share evolutionary trajectories such as gene-by-gene DC, accelerated pseudogenization on neo-X, and pseudogenization pattern. This evolutionary path may be a key for organisms with sex chromosomes to avoid becoming an evolutionary dead-end. Further studies with a broad range of organisms with young sex chromosomes are required to evaluate whether the patterns reported in this study are applicable to other groups of organisms.

## METHODS

### Flies

*D. americana* (strain 15010-0951.03, Millersberg, USA), *D. texana* (strain 15010-1041.00, St. Francisville, USA), and *D. novamexicana* (strain 15010-1031.00, Grand Junction, USA) were obtained from *Drosophila* Species Stock Center (https://www.drosophilaspecies.com/). *D. albomicans* (NG-3, Nago, Japan), *D. nasuta* (strain G-86, Curepipe, Mauritius), and *D. kohkoa* (strain X-145, Kuala Lumpur, Malaysia) have been maintained as living stocks at Tokyo Metropolitan University since the late 1970s.

### Genome and RNA sequencing

For genome sequencing, high molecular weight genomic DNA was extracted from males and females separately. Whole genome shotgun sequencing was performed using the Illumina HiSeq 2500 and PacBio RSII/Sequel platforms (Pacific Biosciences, Menlo Park). PacBio long reads were used to assemble the genome using the Hierarchical Genome Assembly Process (HGAP) 3 or HGAP4 (Chin et al. 2013). Illumina short-reads were then used to correct sequence errors using BWA (Li and Durbin 2009). To obtain the neo-Y assemblies of *D. americana* and *D. albomicans*, we replaced the neo-X sequences with male-specific variants. The genome assemblies of the nine species used in this study were evaluated using BUSCO ver. 4.0.2 software (Simao et al. 2015).

For RNA sequencing, RNA was extracted from third instar larvae, pupae, adults, ovaries, and testes, and a cDNA library from each sample was sequenced using HiSeq 4000 or HiSeq X (Illumina, San Diego). We prepared two biological replicates for each condition to account for the fluctuation among samples. Transcriptome assembly for each species was obtained using HISAT2-2.1.0 (Kim et al. 2019) and StringTie 1.3.6 (Pertea et al. 2015). TransDecoder-5.5.0 (Haas et al. 2013) and Augustus-3.3.2 (Stanke and Waack 2003) were used to annotate coding sequences of expressed and non-expressed genes, respectively. The expression level for each gene was estimated using the STAR 2.7 (Dobin et al. 2013)-RSEM v1.3.1 (Li and Dewey 2011) pipeline.

### Gene classification

To identify orthologs in the three closely-related species (i.e., *D. miranda*–*D. pseudoobscura*–*D. obscura, D. albomicans*–*D. nasuta*–*D. kohkoa*, or *D. americana*–*D. texana*–*D. novamexicana*) and inparalogs for each species, we conducted BLASTP (ver. 2.9.0+) search (Altschul et al. 1997). All identified genes were classified into functional, silenced, disrupted, silenced-and-disrupted, and unclassified genes based on coding sequence integrity and expression level.

See *SUPPLEMENTARY METHODS* for complete methods with detailed descriptions.

## DATA ACCESS

All sequence data reported in this study are available in the DDBJ Sequence Read Archive under the accession numbers DRA007619-26, DRA008052, DRA006582, and DRA004735 (see also Tables S1 and S4 for accession numbers for each run). Genome assemblies for the six *Drosophila* species are also available in the DDBJ under the accession numbers BJEI01000001-BJEI01000834 for *D. albomicans*, BJEH01000001-BJEH01000604 for *D. nasuta*, BJEL01000001-BJEL01000087 for *D. kohkoa*, BJEJ01000001-BJEJ01000435 for *D. americana*, BJEK01000001-BJEK01000199 for *D. texana*, and BJEM01000001-BJEM01000270 for *D. novamexicana* (see also Table S2). Other data are also available upon a request to M.N.

## ACKNOWLEDGMENTS

We thank Yasuko Urabe for her careful experiments. We also thank Kentaro M. and Tanaka, Ryo Yamaguchi for their comments on earlier versions of our manuscripts. We would like to thank the staff of Comparative Genomics Laboratory at National Institute of Genetics for supporting genome sequencing. This work was supported by JSPS KAKENHI Grant Numbers 25711023, 15K14585, 17H05015, 221S0002, and 16H06279 and by the NBRP Genome Information Upgrading Program.

## AUTHOR CONTRIBUTIONS

MN designed the research. MN, KS, SK, and KT prepared the fly materials. MN, YM, and AT conducted DNA sequencing. MN conducted other experiments. MN analyzed the data. MN wrote the draft version of the manuscript. All authors checked the manuscript and approved the research contents.

